# AtlasXbrowser enables spatial multi-omics data analysis through the precise determination of the region of interest

**DOI:** 10.1101/2022.05.11.491526

**Authors:** Joshua Barnett, Jonah Silverman, Molly Wetzel, Poorvi Rao, Noori Sotudeh, Liya Wang

## Abstract

Recent developments in novel spatial sequencing technologies allow for the incorporation of spatial information into high-throughput sequencing assays. One such method, Deterministic Barcoding in Tissue for spatial omics sequencing (DBiT-seq, abbreviated herein as DBiT), utilizes perpendicular microfluidic channels to deliver DNA barcodes across the tissue in a spatially-encoded manner, allowing for sequenced reads to be mapped back onto the 2-D coordinates of the tissue to provide spatial coordinates to cells. DBiT has been the first spatial sequencing technology developed for epigenomic assays beyond transcriptome and proteome. However, despite existing of many open-source software packages for downstream bioinformatics analysis, there is no software available for processing DBiT image data with evenly spaced channels. To facilitate the integration of DBiT spatial and sequenced data, here we proposed a new method to precisely capture the spatial information and further developed **AtlasXbrowser** based on the new method to extract spatial data from the image data.

AtlasXbrowser is a python-based tool with GUI that requires no technical expertise to operate and enables researchers to incorporate brightfield and epifluorescence images of processed tissue samples into downstream bioinformatics analysis tools.

**Availability and implementation:** Freely available at https://github.com/atlasxomics/AtlasXbrowser.

## 1 Introduction

The spatial location and the biological neighborhood of a cell play a critical role in both healthy cellular function (van Vliet *et al*., 2018; Asp *et al*., 2019; Rödelsperger *et al*., 2021) and disease states (de Bruin *et al*., 2014; Maniatis *et al*., 2019; Chen *et al*., 2020). Spatial sequencing offers researchers the exciting prospect of capturing this critical spatial information in their omics analysis (Marx, 2021). For the research community to capitalize on the potential of the technology, performing a spatial sequencing assay must be easy, accurate, and reproducible. Achieving these goals entails the development of tools used to facilitate the workflows themselves. Here we highlight **AtlasXbrowser**, a free open source python-based GUI, created to streamline the image processing necessary to perform any DBiT assay (Liu *et al*., 2020; Deng *et al*., 2022). In particular, AtlasXbrowser makes it easy to localize transcriptome, protein, or epigenome signals on a tissue section.

The DBiT method involves the sequential flow of unique spatial barcodes through microfluidic channels. The *x* and *y* barcodes are ligated to target biomolecules. Each intersection area between a microfluidic row and column (*x* and *y*) generates an element of tissue defined herein as a TIXEL™ unit (abbreviation for tissue pixel). Processing the library of spatially encoded target biomolecules through next-generation sequencing (NGS) and feeding the raw reads through the bioinformatics workflow permits reconstruction of a mosaic of TIXEL units. Following the application of both microfluidic chips (A chip with horizontal channels and B chip with vertical channels) to the tissue, AtlasXbrowser guides the user through the process of locating the region of interest (ROI), defined as the pixels of the micrograph corresponding to the location of the TIXEL mosaic. This process also permits the manual or automatic classification of each TIXEL unit as being either on or off tissue.

AtlasXbrowser encapsulates the numerous advances made in the DBiT protocol since its inception. Whereas previously the ROI was determined by an eyeball estimation of the overlapping marks on the tissue from the A and B chip channels, the advancement of flowing bovine serum albumin (BSA) mixed with a fluorescent dye through the outmost channels of the chips in combination with AtlasXbrowser’s ROI designation functionality allows for a much more precise ROI designation and thus the ability to more accurately determine TIXEL unit coordinates. Further, while previously global thresholding was used to binarize the tissue images, AtlasXbrowser incorporates adaptive thresholding (Bradski, 2000) to allow for a more flexible pixel classification system, accounting for variation in inhomogeneous tissue, lighting, and contrast that may be present across various regions of the image. Once the binarized image has been created, AtlasXbrowser creates the infrastructure for researchers to identify on and off tissue TIXEL units both when initially processing the image, and by loading the annotated images back into AltasXbrowser in combination with sequencing data. Additionally, AtlasXbrowser does not assume the A and B channels are perpendicular to each other as previously which makes the placement of A and B chips more flexible. Finally, AtlasXbrowser has standardized the output of DBiT image data, creating a “Spatial” folder, containing the output of the image processing in the 10x Visium image data format that can be used in most bioinformatic packages for incorporating spatial information. The usage of AtlasXbrowser is agnostic to the assay being performed, and thus serves as one tool which can be used for DBiT experiments for multi-omics.

## 2 Methods

### 2.1 Implementation

AtlasXbrowser is written in Python 3.7 and uses the cross-platform Tkinter library (Lundh, 1999) to build a user-friendly graphical interface. Image operations including adaptive thresholding are built with the python OpenCV library (Bradski, 2000). Detailed documentation is provided with a guided example: https://docs.atlasxomics.com/projects/AtlasXbrowser/.

### 2.2 Data

#### 2.2.1 DBiT-seq Assay

Following the spatial Cut&Tag protocol (Deng *et al*., 2022), a DBiT assay is performed using the H3K27ac antibody to reveal the spatial map of potentially upregulated genes, as highlighted in Figure 1.

**Figure 1:**
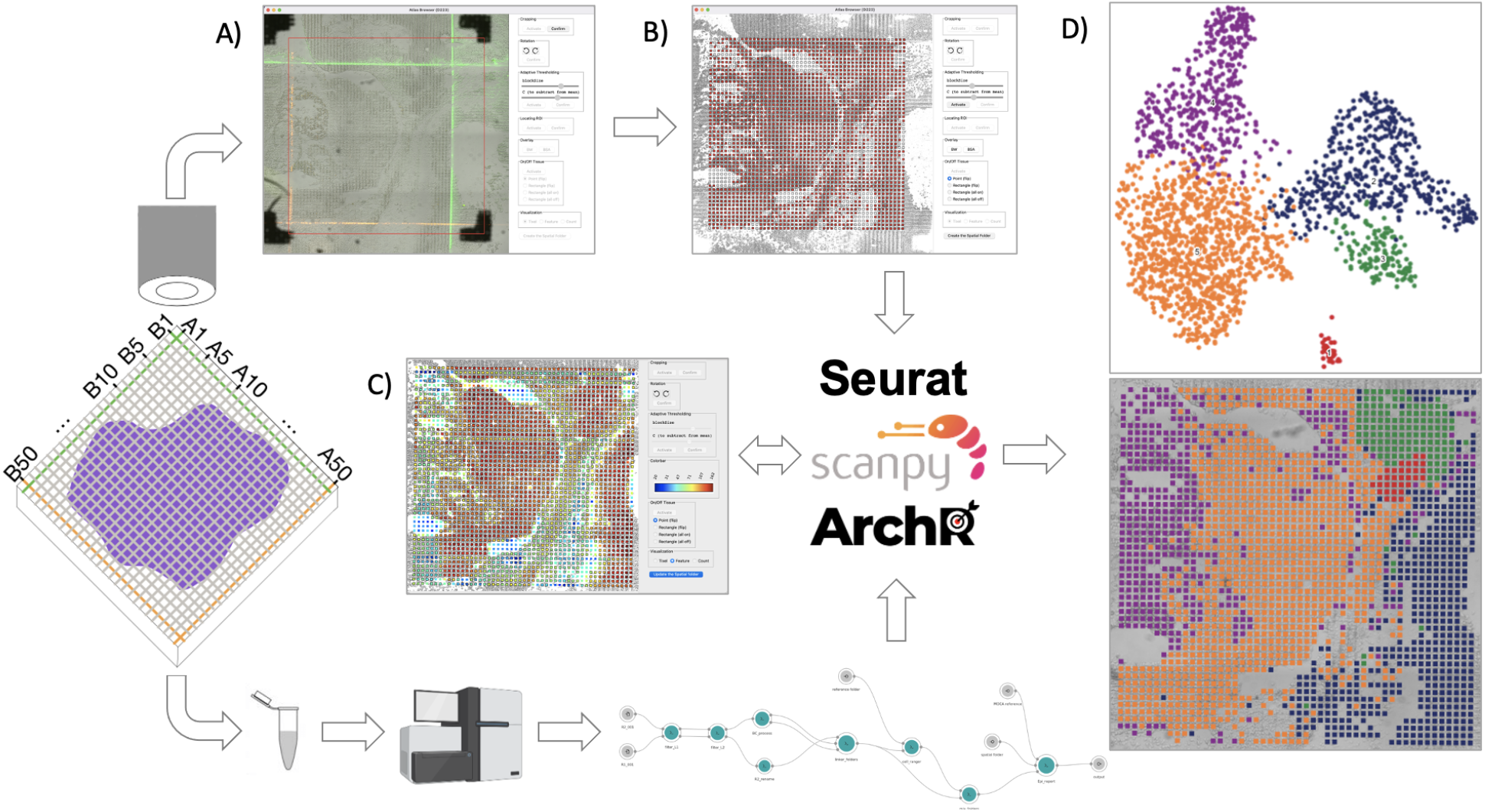
Overview of how AtlasXbrowser facilitates spatial data analysis. After removing the B chip, images are captured with a microscope and processed along with user input to generate the locations of the intersection points between channels in the two chips. The tissue section is lysed to prepare a sequencing library which is sequenced and processed with a bioinformatics workflow to generate counts. Then sequencing counts and spatial data are combined for spatial data analysis. A.) Localizing the ROI by fitting a draggable and resizable quadrilateral to the outside boundary of the BSA lines. B) After binarization with adjustable adaptive thresholding, interactively labeling TIXEL units as on or off tissue based on the intensity of brightfield pixels. C) Interactively adjusting on/off tissue TIXEL units with fragment (epigenome) or UMI/gene (transcriptome) counts derived from sequencing data and displayed with colormap. D) Umap (top) and spatial (bottom) plot by Seurat/Scanpy/ArchR after combining sequencing data with spatial data and correcting for on/off tissue classifications from B and C.

#### 2.2.2 Brightfield and Fluorescence Imaging

To precisely determine the position of the ROI, the original DBiT protocol (Su *et al*., 2021) was modified by flowing both barcode and a solution of bovine serum albumin (BSA) mixed with a fluorescent dye in the first and last channels of both A and B chips. The dye left on the tissue marks the outside boundary of the ROI when the image (so-called BSA image) is taken using the microscope’s fluorescent channel. With the outside boundary marked on the BSA image, AtlasXbrowser calculates the precise location of each TIXEL observation with the assumption that all the microfluidic channels are evenly spaced.

To ensure the image is in the correct orientation, green dye (Fluorescein) is used in the first channel of both the A chip (top channel) and B chip (rightmost channel) while red dye (ROX) is used in the last channel of both A chip (bottom channel) and B chip (leftmost channel). With the correct orientation, the BSA image should have the bottom and left boundaries of the ROI in red while the top and right boundaries of the ROI in green (as shown in Figure 1). Similar to the DBiT protocol, a brightfield image, a so-called postB image, is also saved after removing the B chip.

### 2.3 Image Analysis with AtlasXbrowser

Following the creation of the BSA and postB images, as shown in Figure 1A, the images are loaded into AtlasXbrowser, and the relevant metadata pertaining to the experiment ID is inputted. AtlasXbrowser then guides the user through the process of cropping the image to remove redundant regions, as well as rotating the image as desired. Then using the fluorescent dye as a cue marking the outside boundary of the outmost channels of the ROI, the user identifies the coordinates of the four corners of the ROI through AtlasXbrowser. This information, in conjunction with the knowledge of evenly spaced TIXEL units, allows for the determination of each TIXEL unit’s coordinate location.

Then the tissue image is binarized, converting each pixel in the image into being either black or white. This is done in an effort to determine which regions of the image correspond to biological tissue, and which do not. Then, based on the proportion of black pixels with each TIXEL unit, AtlasXbrowser makes predictions about which TIXEL units are on or off tissue. Following this, the user is able to make any manual adjustments to the designation of on or off tissue for each TIXEL unit or a group of TIXEL units (Figure 1B). Once the TIXEL units have been classified as on vs. off tissue, Altasbrowser generates a “Spatial” folder. This folder contains the cropped and rotated postB and BSA images, a metadata file, as well as all 10x Visium standardized files with information pertaining to TIXEL unit coordinate location with barcode identifiers, on and off tissue state of all TIXEL units.

### 2.4 Spatial Bioinformatics Analysis

In conjunction with the sequencing data for the tissue, the spatial data generated by AtlasXbrowser can be inputted directly into bioinformatic analysis tools, such as Seurat (Hao *et* al., 2021), Scanpy (Wolf *et al*., 2018), or ArchR (Granja *et al*., 2021). Examples of this are shown in Figure 1D where UMAP and Spatial plots are shown by combining sequencing and spatial data. Note that the bioinformatics analysis results, if added to the “Spatial” folder, can also be loaded back into the AtlasXbrowser to further refine on/off tissue designations, as shown in Figure 1C.

## 3 Discussion

DBiT offers the research community the ability to perform spatial transcriptome (Liu *et al*., 2020), ATAC (Deng *et al*., 2021), Cut&Tag sequencing (Deng *et al*., 2022), as well as co-mapping of transcriptome and protein (Liu *et al*., 2022), or transcriptome and ATAC (Peng *et al*., 2022) while preserving the tissue spatial information. AtlasXbrowser allows for an assay agnostic image processing GUI that can be used for all DBiT assays. Recent advances in the DBiT workflow such as the flowing of fluorescently tagged BSA solution through the outmost channels of the A and B chips are capitalized in the AtlasXbrowser process, and more so AtlasXbrowser facilities improvements in image thresholding by employing adaptive thresholding when binarizing the challenging brightfield images, as well as allowing for the user-specified designation of TIXEL units as on and off tissue, and produces a standardized output in a “Spatial” folder ready for bioinformatic analysis. AtlasXbrowser aims to make DBiT image processing easy, accurate, and reproducible, allowing the research community to more effectively reap the rich insights offered by spatial sequencing and to combine DBiT data with other assays through the precise spatial information.

As the first technology being commercialized for spatial epigenome assays, DBiT does not need sophisticated machines or equipments and thus can be executed in any laboratories with access to a microscope. Given that most of the bioinformatics packages for spatial data analysis are readily accessible, we anticipate that AtlasXbrowser will fill the gap in DBiT data analysis and in turn make DBiT widely applicable to any spatial multi-omics studies.

## Notes

### Competing Interest Statement

The authors have declared no competing interest.

### Summary of Updates

Updated author list

